# The mechanism underlying transient weakness in myotonia congenita

**DOI:** 10.1101/2020.12.23.424129

**Authors:** Jessica H Myers, Kirsten Denman, Chris DuPont, Ahmed A Hawash, Kevin R Novak, Andrew Koesters, Manfred Grabner, Anamika Dayal, Andrew A Voss, Mark M Rich

**Author notes:** Please address all correspondence to: Mark Rich.

## Abstract

In addition to the hallmark muscle stiffness, patients with recessive myotonia congenita (Becker disease) experience debilitating bouts of transient weakness that remain poorly understood despite years of study. We made intracellular recordings from muscle of both genetic and pharmacologic mouse models of Becker disease to identify the mechanism underlying transient weakness. Our recordings reveal transient depolarizations (plateau potentials) of the membrane potential to −25 to −35 mV in the genetic and pharmacologic models of Becker disease. Both Na^+^ and Ca^2+^ currents contribute to plateau potentials. Na^+^ persistent inward current (NaPIC) through Naγ1.4 channels is the key trigger of plateau potentials and current through Ca_v_1.1 Ca^2+^ channels contributes to the duration of the plateau. Inhibiting NaPIC with ranolazine prevents the development of plateau potentials and eliminates transient weakness *in vivo.* These data suggest that targeting NaPIC may be an effective treatment to prevent transient weakness in myotonia congenita.

**Impact Statement:** Transient weakness in myotonia congenita is caused by depolarization secondary to activation of persistent Na^+^ current in skeletal muscle.

## Introduction

Myotonia congenita is one of the non-dystrophic muscle channelopathies. It is caused by loss-of-function mutations affecting the muscle chloride channel (ClC-1) (Lipicky et al., 1971; Steinmeyer et al., 1991; Koch et al., 1992). Patients with recessive myotonia congenita (Becker disease) experience muscle stiffness due to hyperexcitability (Lehmann-Horn et al., 2008; Trivedi et al., 2014; Cannon, 2015) as well as transient weakness due to unknown factors (Ricker et al., 1978; Rudel et al., 1988; Zwarts and van Weerden, 1989; Deymeer et al., 1998; Trivedi et al., 2013). Some patients with Becker disease report transient weakness in arm muscles as a greater impediment than muscle stiffness (Rudel et al., 1988). This weakness can last up to 90 seconds and is brought on by exertion following rest (Ricker et al., 1978; Rudel et al., 1988; Zwarts and van Weerden, 1989; Deymeer et al., 1998).

The mechanism underlying transient weakness in Becker disease has remained unknown since its initial description close to 50 years ago (Ricker and Meinck, 1972). There appears to be loss of muscle excitability, as weakness is accompanied by a drop in compound muscle action potential amplitude (CMAP) during repetitive stimulation (Ricker and Meinck, 1972; Brown, 1974; Aminoff et al., 1977; Deymeer et al., 1998; Drost et al., 2001; Modoni et al., 2011). This drop in CMAP is associated with reduction in muscle fiber conduction velocity, which has been proposed to progress to depolarization block (Zwarts and van Weerden, 1989). What has remained unclear, and perhaps counterintuitive, is why a loss-of-function mutation of the muscle ClC-1 channels in myotonia congenita (Lipicky et al., 1971; Steinmeyer et al., 1991; Koch et al., 1992) leads to transient loss of excitability. The primary defect caused by loss of ClC-1 current is hyperexcitability of muscle, which causes myotonia.

We established that a ClC-1 homozygous null (ClC^adr^) mouse model of Becker disease has transient weakness *in vivo*, mimicking the condition in human patients. Intracellular recording in both ClC^adr^ muscle and a pharmacologic model of Becker disease (due to block of ClC-1 with 9-AC) have elucidated a novel phenomenon: transient depolarizations to voltages between −25 to −35 mV, lasting many seconds, which we termed “plateau potentials.” Blocking Na^+^ persistent inward current (NaPIC) with ranolazine (a piperazine derivative) prevented both development of plateau potentials and transient weakness. We conclude that NaPIC plays a central role in the development of plateau potentials, which are the mechanism underlying transient weakness in Becker disease.

## Results

### Identification of transient weakness in mice with myotonia congenita

To study *in situ* isometric motor performance in the *Clcn1*^adr-mto2J^ (ClC^adr^) mouse model of recessive myotonia congenita (Becker disease), a muscle force preparation that we used previously was employed (Dupont et al., 2019; Wang et al., 2020). Mice were anesthetized via isoflurane inhalation and the distal tendon of the triceps surae (gastrocnemius, plantaris, and soleus muscles) was dissected free and attached to a force transduction motor; then the sciatic nerve was stimulated with 45 pulses delivered at 100 Hz. In unaffected littermates, there was no myotonia following 45 pulses at 100 Hz, such that relaxation was immediate (Fig 1A). In ClC^adr^ mice, stimulation with 45 pulses caused full fusion of force, but relaxation was slowed, due to the presence of myotonia (Fig 1C).

**Fig 1:**
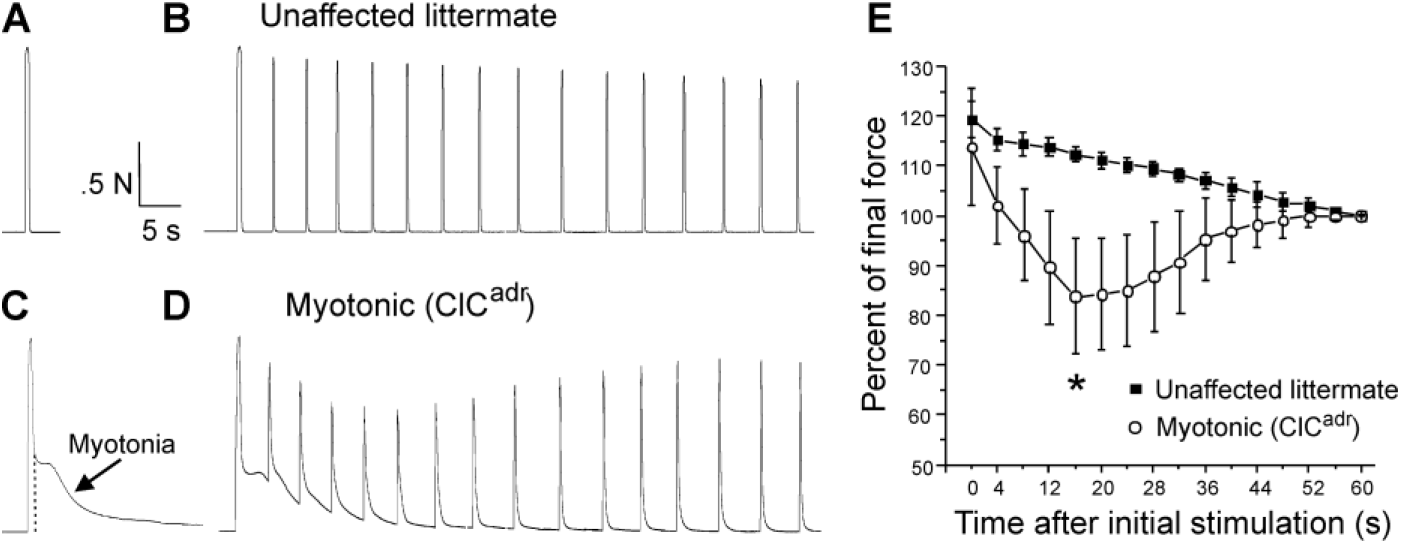
Transient weakness in the ClC^adr^ mouse model of recessive myotonia. A) Shown is a force trace from a triceps surae muscle group of an unaffected littermate in response to stimulation of the sciatic nerve with 45 pulses at 100 Hz. B) The force trace generated by following the initial stimulus with 15 pulses at 100 Hz delivered every 4 s. C) In a ClC^adr^ mouse, following the 45 pulses at 100 Hz (indicated by the dotted vertical line) there was continued force generation secondary to myotonia. D) With stimulation every 4 s transient weakness was revealed. E) Plot of the force normalized to force at 60 s in unaffected littermates and ClC^adr^ mice. Transient reduction in force was present in ClC^adr^ mice (*, p = .00015 vs unaffected littermates at 16s, t-test, 95% confidence interval 110 to 114 vs 75 to 93). n = 5 unaffected littermates and n = 9 ClC^adr^ mice. Error bars represent ± SD.

To determine whether transient weakness was present, the sciatic nerve was additionally stimulated with 15 pulses at 100 Hz every 4 seconds for 1 minute. In unaffected littermates, this caused stable force production with a mild, gradual reduction that was likely due to fatigue (Fig 1B). In myotonic mice, the same stimulation protocol revealed transient weakness, as force fell over the first 10 to 15s and then recovered (Fig 1D). To avoid inclusion of fatigue in measurement of transient weakness, we normalized to force at the end of the 1 minute of intermittent stimulation. The plot of the mean normalized force revealed transient weakness, which peaked in severity 15 to 20s after the initial stimulation and resolved within 1 minute (Fig 1E). These data indicate transient weakness is present in ClC^adr^ mice.

### Characterization of plateau potentials in genetic and pharmacologic models of myotonia congenita

In myotonic patients, transient weakness is paralleled by a drop in compound muscle action potential amplitude (CMAP) (Ricker and Meinck, 1972; Modoni et al., 2011). This finding suggests weakness is due to loss of excitability. To look for inexcitability of ClC^adr^ muscle, intracellular current clamp recordings were performed. In both unaffected littermates and ClC^adr^ mice, stimulation with a 200-ms injection of depolarizing current triggered repetitive firing of action potentials during the stimulus. In muscle from unaffected littermates, the firing ceased as soon as the stimulus was terminated (Fig 2A). In muscle from ClC^adr^ mice, there was myotonia (continued firing of action potentials following termination of the stimulus) in 100% of fibers (Fig 2B). The myotonia often persisted for many seconds. While many runs of myotonia ended with repolarization to the resting membrane potential (Fig 2B), in other instances, myotonia terminated with development of depolarizations lasting 5 to >100s to a membrane potential near −30 mV (Fig 2C, D). During the depolarizations there was gradual repolarization of the membrane potential, which was followed by sudden repolarization back to the resting potential. In some cases, the sudden repolarization was preceded by development of oscillations in the membrane potential (Fig 2D). The depolarizations occurred in 62% of ClC^adr^ muscle fibers (n = 42 fibers from 10 mice). Depolarizations lasting less than 1s to a membrane potential close to −60 mV have been described in a toxin-induced model of hyperkalemic periodic paralysis and were termed plateau depolarizations (Cannon and Corey, 1993). While the depolarizations we identified could be due to similar mechanisms as those described in hyperkalemic periodic paralysis, they seemed more similar to transient depolarizations in spinal motor neurons, which can last many seconds, and have been termed plateau potentials (Alaburda et al., 2002; Heckman and Enoka, 2012; Hounsgaard, 2017). We thus chose the term plateau potentials to describe them.

**Fig 2.**
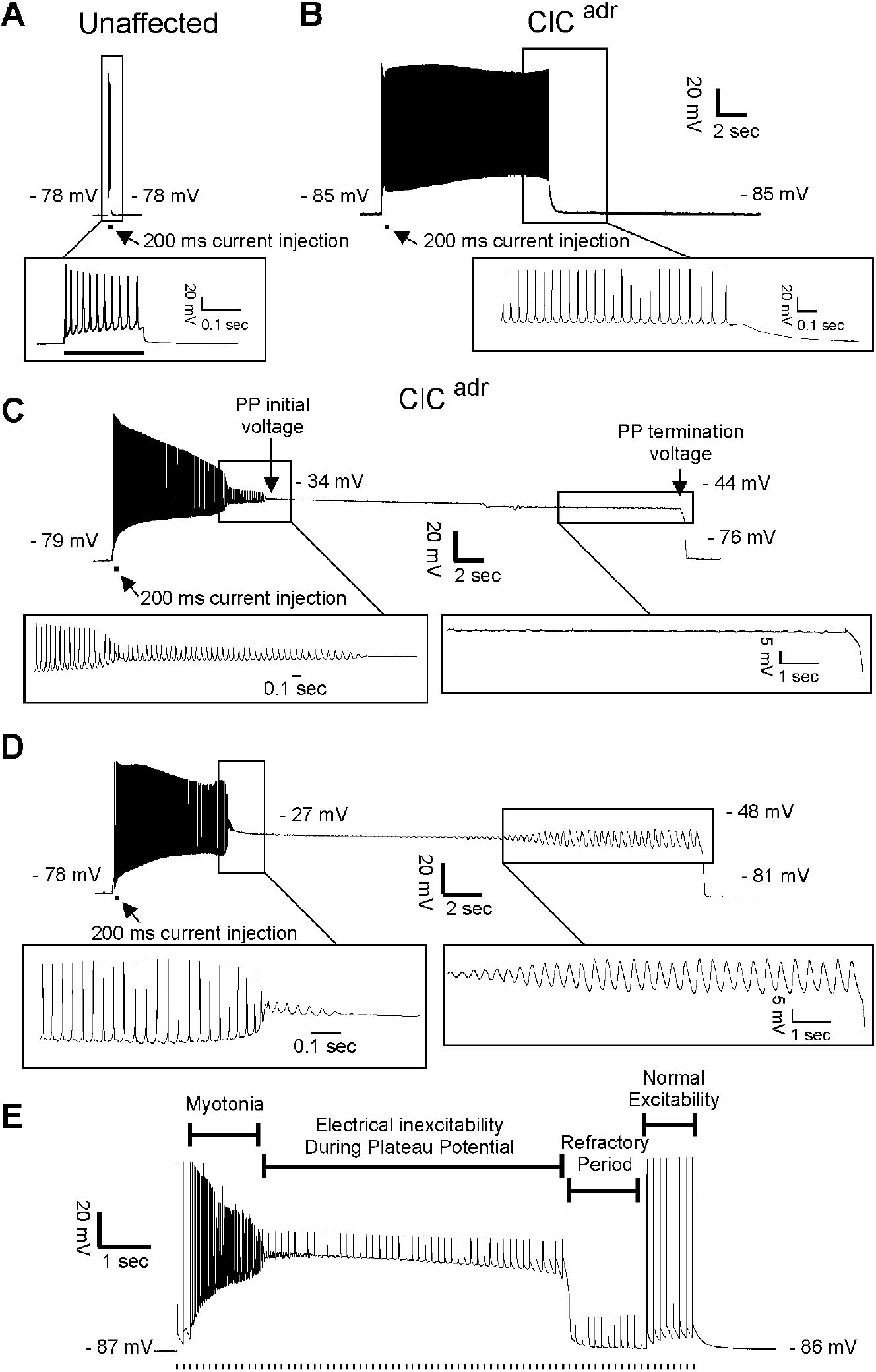
Plateau potentials in ClC^adr^ muscle. For A through D, the insets show portions of the traces on an expanded time base. A) The response of muscle from an unaffected littermate to injection of 200ms of depolarizing current (horizontal bar below the voltage trace); note that action potentials stop when current stops. B-D) traces of myotonia triggered by a 200-ms injection of depolarizing current from 3 different ClC^adr^ muscle fibers. The following membrane potentials are identified in C and D: membrane potential prior to stimulation, initial membrane potential during the plateau potential, membrane potential prior to the termination of the plateau potential, and membrane potential following repolarization. E) Development of a plateau potential during repetitive stimulation at 8 Hz (stimuli represented by vertical hash marks under the recording).

To determine whether plateau potentials cause inexcitability, ClC^adr^ muscle fibers were stimulated during the plateau phase. During, and immediately following plateau potentials, action potential generation in response to current injection failed (Fig 2E). At later times following repolarization, action potential generation was again possible. The inexcitability at earlier times following repolarization is likely due to slow inactivation of Na channels following the many-second depolarization (Ruff, 1996, 1999; Rich and Pinter, 2003). These data are consistent with the possibility that plateau potentials and the resultant inexcitability of muscle are the mechanism underlying transient weakness in myotonia congenita.

In ClC^adr^ muscle, ClC-1 chloride conductance has been absent throughout development such that plateau potentials could be a compensatory response to muscle hyperexcitability. To test this possibility, we acutely blocked ClC-1 chloride channels in muscle from unaffected littermates with 100 μM 9-anthracene carboxylic acid (9-AC). This dose of 9-AC blocks more than 95% of ClC-1 chloride channels in skeletal muscle (Palade and Barchi, 1977) and has been used to model myotonia congenita both *in vitro* and *in vivo* (van Lunteren et al., 2011; Desaphy et al., 2013; Desaphy et al., 2014; Skov et al., 2015). Acute blocking of ClC-1 triggered myotonia and plateau potentials in 94% of fibers (Fig 3A, n= 53 fibers from 8 mice). These data strongly suggest that the ion channels responsible for development of plateau potentials are present in wild-type skeletal muscle.

**Fig 3:**
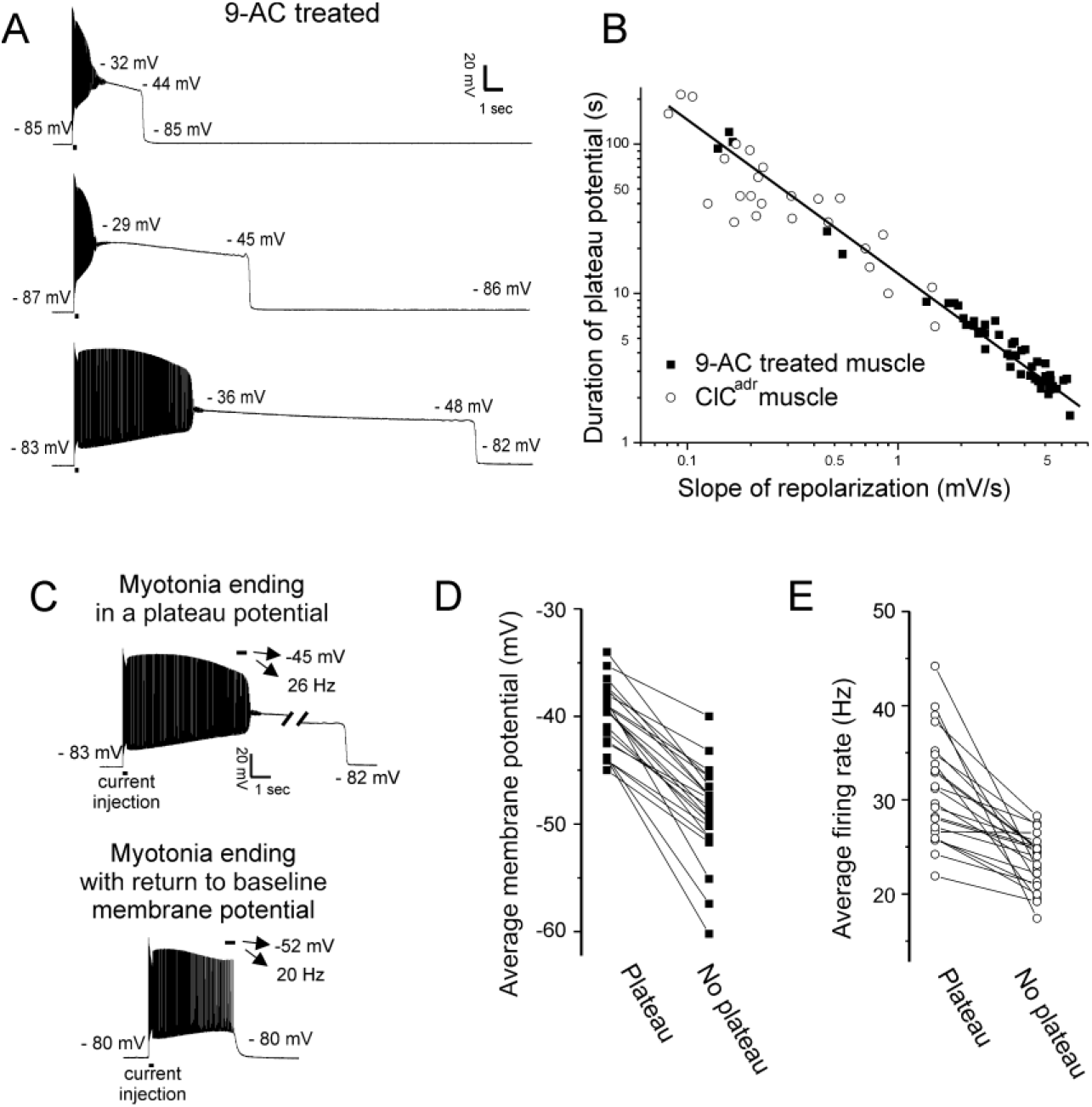
Characterization of plateau potentials. A) Three examples of plateau potentials of different duration in 9-AC treated muscle. The 200-ms current injection is indicated by a horizontal bar underneath the trace. Indicated on each trace is the membrane potential at the start and end of the plateau potential. B) The duration of plateau potentials in the ClC^adr^ and 9-AC models of myotonia plotted against the rate of repolarization during the plateau potential. A linear fit was performed on the log-log plot with an R^2^ value of 0.90 and a slope of −0.97. C) Shown are two runs of myotonia from the same muscle fiber, one ending in a plateau potential and one ending with repolarization. The mean membrane potential and firing rate in the final 500ms of myotonia (horizontal bar) are indicated by the arrows above each trace. D) Plot of the average membrane potential during the final 500 ms of myotonia for 22 fibers in which there were both a run of myotonia ending in a plateau potential and a run ending with repolarization (No plateau potential). E) Plot of the average firing rate during the final 500 ms of myotonia for the same 22 fibers.

In contrast to the lack of variability in voltage at onset and termination of plateau potentials, the duration in both the ClC^adr^ and 9-AC treated models of myotonia was highly variable (Fig 3B, Table 1, variance = 3216s for ClC^adr^ and 623s for 9-AC treated duration). The reason for the high variance of duration was that the rate of repolarization during the plateau potential varied by more than 100-fold (Fig 3B, Table 1). It was generally not possible to record multiple plateau potentials in individual ClC^adr^ fibers due to the long median duration. However, it was possible to record multiple plateau potentials within individual fibers of 9-AC treated muscle, as most lasted only a few seconds. Within individual fibers, the slope of repolarization of plateau potentials was less variable such that duration was relatively constant with a mean variance of 0.5s ± 0.7s (n = 17 9-AC treated fibers in which 4 or more plateau potentials were recorded).

**Table 1:** Plateau potential parameters in ClC^adr^ and 9-AC treated unaffected muscle

|  | Pre-myotonia Vm (mV) | Initial PP Vm (mV) | PP repol slope (mV/s) | Final PP Vm (mV) | Max repol rate (mV/s) | Post PP Vm (mV) | Duration of PP (s) |
| --- | --- | --- | --- | --- | --- | --- | --- |
| ClC<sup>adr</sup> | -79.3 ± 2.9<br>(-79.6) | -29.7 ± 3.6<br>(-29.6) | -1.3 ± 4.4<br>(-0.2) | -43.0 ± 5.4<br>(-43.3) | 142 ± 41<br>(138) | -75.7 ± 4.3<br>(-76.0) | 57.6 ± 56.7<br>(42.2) |
| 9-AC | -81.5 ± 2.6<br>(-82.4) | -31.2 ± 3.1<br>(-31.4) | -3.4 ± 1.8*<br>(-3.5) | -45.2 ± 2.3<br>(-45.1) | 197 ± 44**<br>(192) | -80.8 ± 2.9<br>(-81.6) | 11.7 ± 25.0***<br>(4.1) |

Shown are the mean value ± the standard deviation for parameters of plateau potentials in ClC<sup>adr</sup> fibers and 9-AC treated fibers. Below the mean value is the median value in parenthesis. n = 26 fibers with plateau potentials from 10 ClC<sup>adr</sup> mice and 50 fibers with plateau potentials from 8 unaffected muscles treated with 9-AC. Vm is membrane potential, repol is repolarization. Pre-myotonia Vm is the resting membrane potential prior to stimulation. Initial PP Vm is the membrane potential at the beginning of the plateau potential. Final PP Vm is the membrane potential prior to the sudden repolarization terminating the plateau potential. Max repol rate is the maximum rate of repolarization during the sudden repolarization. Post PP Vm is the membrane potential following termination of the plateau potential. \* indicates p = .017 of log transformed data (t-test), \*\* p = 0.001 (t-test), \*\*\* p = 4.5 x 10<sup>-4</sup> of log transformed data (t-test).

In both ClC^adr^ muscle and 9-AC treated muscle, plateau potentials did not occur following every run of myotonia (Fig 3C). To determine why some runs of myotonia ended in plateau potentials while others did not, we compared the mean voltage at the end of runs of myotonia that produced plateau potentials vs. the mean voltage at the end of runs that did not produce plateau potentials, in 9-AC treated fibers. There was a strong correlation between the mean membrane potential during the final 500-ms of runs of myotonia and development of plateau potentials. In 22/22 fibers, the mean membrane potential was more depolarized in runs of myotonia that produced plateau potentials (mean = −40.1 ± 3.1 mV vs −49.0 ± 4.5 mV, p = 5 x 10^-11^, paired t-test, Fig 3D). This suggested that a voltage-dependent current might be involved.

However, another feature of myotonia that determines whether a plateau potential is triggered is the firing rate, which might correlate with elevation of intracellular Ca^2+^. Thus we examined and found a strong correlation between the firing rate of myotonia runs and subsequent development of plateau potentials: In 21/22 fibers, the mean firing rate was higher for runs of myotonia ending in plateau potentials (mean = 31.2 ± 5.7 Hz vs 23.4 ± 2.9, p = 7 x 10^-6^, paired t-test, Fig 3E). Thus, while voltage-dependent channels might be involved, this indicated possible participation of channels gated by other factors, such as elevation of intracellular Ca^2+^.

### Ca^2+^ and Na^+^ currents contribute to generation of plateau potentials

In spinal motor neurons, L-type Ca^2+^ channels play a central role in generation of plateau potentials (Alaburda et al., 2002; Heckman and Enoka, 2012; Hounsgaard, 2017). To determine whether Ca^2+^ current through skeletal muscle L-type channel (Ca_v_1.1) triggers plateau potentials, we performed recordings on skeletal muscle fibers from a mouse model (*nc*DHPR) in which the pore region of Ca_v_1.1 carries a point mutation leading to ablation of inward Ca^2+^ current (Dayal et al., 2017). We used voltage clamp of FDB/IO fibers to verify the absence of inward currents in *nc*DHPR mice. In wild-type littermate controls, ramp depolarization following block of Na^+^, K^+^, and Cl^-^ channels triggered a large, inward Ca^2+^ current, which began to activate at −15.1 ± 6.9 mV (n= 4 fibers) with a mean amplitude of 165 ± 46 nA (Fig 4 A, B). Ramp depolarization triggered no inward Ca^2+^ current in *nc*DHPR muscle fibers (mean = 0 ± 0 nA, n = 10 fibers (Fig 4C).

**Fig 4:**
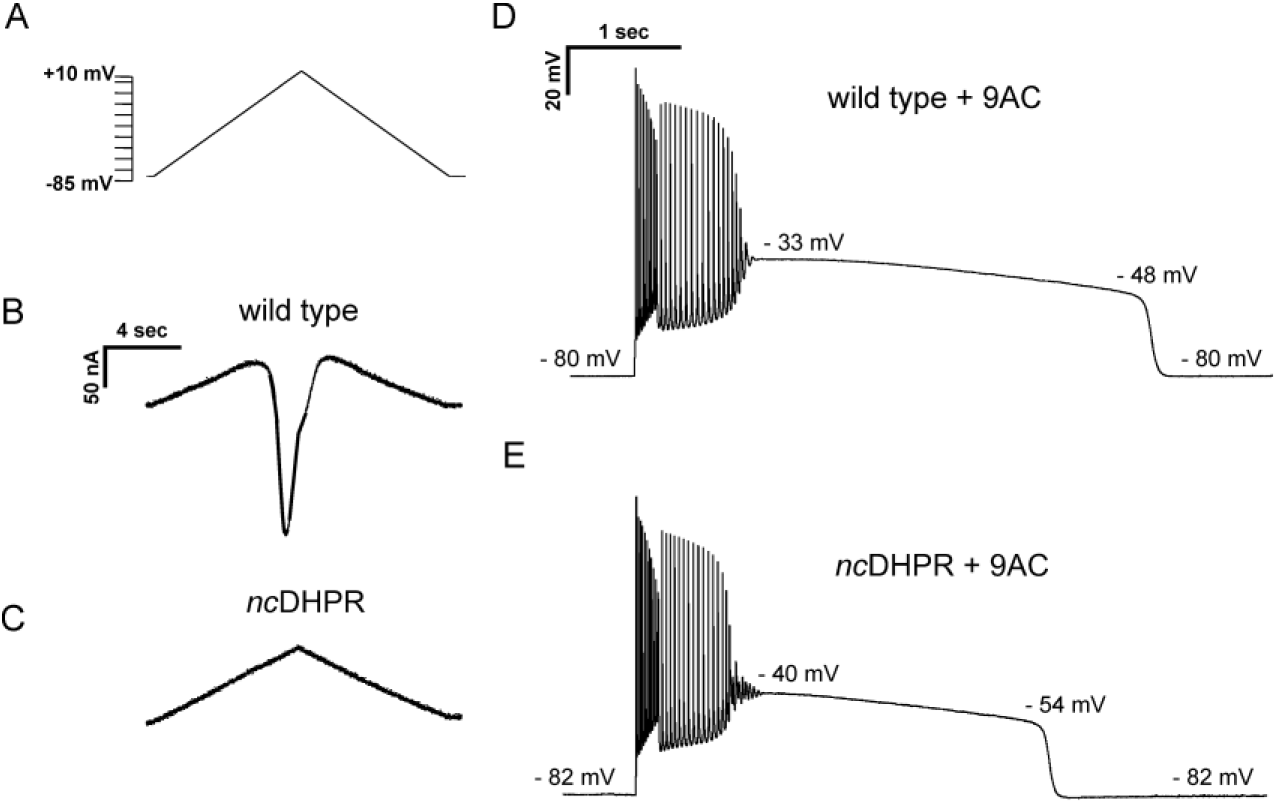
Current flow through Ca_v_1.1 does not initiate, but helps to sustain, plateau potentials in 9-AC treated muscle. A) The voltage protocol applied to FDB/IO fibers consisted of a ramp depolarization from −85 mV to +10 mV applied over 8 s. B) A large inward current is present in wild-type muscle when Na^+^, K^+^, and Cl^-^ currents were blocked. C) *nc*DHPR muscle fibers confirm the absence of inward Ca^2+^ current as there is only the linear change in current due to the changing command potential. D) A plateau potential in a 9-AC treated wild-type muscle fiber. E) A plateau potential in a 9-AC treated *nc*DHPR fiber.

To determine whether current flow through Ca_v_1.1 contributes to generation of plateau potentials, current clamp recordings were performed. Application of 100 μM 9-AC led to development of plateau potentials in 41/49 fibers from 4 wild-type mice and in 47/49 fibers from 3 *nc*DHPR mice (Fig 4D and E). Analysis of the characteristics of plateau potentials suggests that Ca^2+^ current flow through Ca_v_1.1 channels may influence the duration of the plateau. While there was no difference in the beginning or ending voltages of plateau potentials, the duration was shorter in *nc*DHPR muscle (Fig 4D and E, Table 2). The cause of the shorter plateau potential was an increase in the rate of repolarization (Table 2). These data suggest that Ca^2+^ influx through Ca_v_1.1 does not play a role in initiation of plateau potentials, but is involved in sustaining them.

**Table 2:** Plateau potential parameters in 9-AC treated wild-type and *nc*DHPR muscle

|  | Pre-myotonia<br>Vm (mV) | Initial plateau<br>Vm (mV) | PP repol<br>slope (mV/s) | Final PP Vm<br>(mV) | Max repol<br>rate (mV/s) | Post PP Vm<br>(mV) | Duration of PP<br>(s) |
| --- | --- | --- | --- | --- | --- | --- | --- |
| wild-type | $-79.3 \pm 0.9$<br>(-79.5) | $-36.0 \pm 1.6$<br>(-36.0) | $-3.3 \pm 0.7$<br>(-3.1) | $-48.1 \pm 1.2$<br>(-48.5) | $-125 \pm 6$<br>(125) | $-79.5 \pm 0.4$<br>(-79.7) | $3.9 \pm 0.7$<br>(3.8) |
| *ncDHPR* | $-78.9 \pm 0.6$<br>(-79.0) | $-37.2 \pm 0.8$<br>(-37.1) | $-5.4 \pm 0.6^*$<br>(-5.3) | $-49.2 \pm 1.1$<br>(-49.7) | $-134 \pm 10$<br>(138) | $-79 \pm 0.8$<br>(-79.3) | $2.4 \pm 0.3^*$<br>(2.3) |

Shown are the mean value and the standard deviation for parameters of plateau potentials in 9-AC treated wild-type and *ncDHPR* fibers. Below the mean value in parentheses is the median value. $n = 41$ fibers from 4 wild-type mice and $n = 47$ fibers from 3 *ncDHPR* mice. Vm is membrane potential, repol is repolarization. Pre-myotonia Vm is the resting membrane potential prior to stimulation. Initial PP Vm is the membrane potential at the beginning of the plateau potential. Final PP Vm is the membrane potential prior to the sudden repolarization terminating the plateau potential. Max repol rate is the maximum rate of repolarization during the sudden repolarization. Post PP Vm is the membrane potential following termination of the plateau potential. \* indicates $p = 0.01$ for both significant differences (t-test).

The Na_v_1.4-mediated Na^+^ current responsible for generation of action potentials in skeletal muscle inactivates within ms of depolarization, making it highly unlikely that it contributes to development of plateau potentials. However, a Na^+^ current lacking fast inactivation (Na^+^ persistent inward current, NaPIC) was found to be involved in the generation of plateau depolarizations in muscle in a toxin model of hyperkalemic periodic paralysis (Cannon and Corey, 1993). We recently determined that NaPIC contributes to repetitive firing occurring during myotonia (Hawash et al., 2017; Metzger et al., 2020). NaPIC is present in normal skeletal muscle and likely derives from a small subset of Na_v_1.4 channels that are in a conformation that differs from fast-inactivating channels (Patlak and Ortiz, 1986; Gage et al., 1989). We term the Na^+^ channels responsible for action potentials “fast-inactivating Na^+^ channels” and Na^+^ channels lacking fast inactivation “NaPIC”.

To determine whether NaPIC might play a role in generation of plateau potentials, we applied ranolazine to 9-AC treated muscle. Ranolazine has been found to preferentially block NaPIC in brain, heart, peripheral nerve, and skeletal muscle (El-Bizri et al., 2011; Kahlig et al., 2014). We previously found that ranolazine was effective in eliminating myotonia by blocking NaPIC while sparing enough fast-inactivating Na^+^ channels to allow for repetitive firing of action potentials triggered by current injection (Novak et al., 2015; Hawash et al., 2017). As shown in Fig 5A, when myotonia was triggered by treatment of muscle with 9-AC, 94% of fibers (n= 53 fibers from 8 mice) developed plateau potentials in response to a 200-ms injection of depolarizing current. Following treatment with 40 μM ranolazine, 0/22 fibers from 3 mice developed plateau potentials after 200-ms current injection (Fig 5A, p < .01 vs untreated). These data were consistent with NaPIC playing a role in development of plateau potentials.

**Fig 5:**
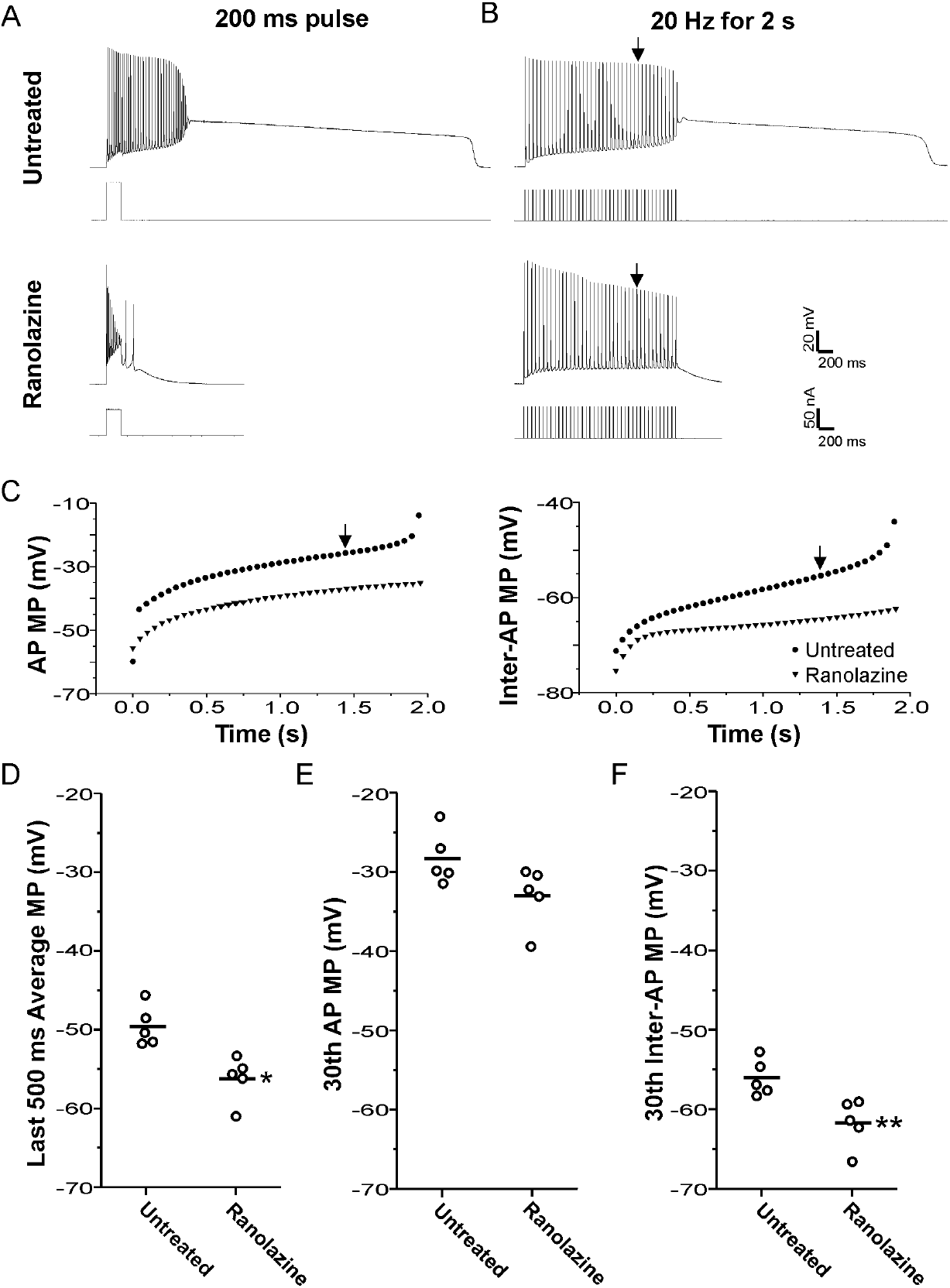
Block of plateau potentials by ranolazine in 9-AC treated muscle is associated with lessening of depolarization during repetitive firing. A) Examples of the response to a 200-ms injection of depolarizing current at baseline and following treatment with 40 μM ranolazine. B) Examples of the response to 2s of stimulation at 20 Hz. The vertical arrow points to the 30^th^ action potential in each trace, which was used for analysis of differences between ranolazine untreated and ranolazine treated muscle (D-F). C) Plots of the membrane potential during the 8 ms encompassing the action potential and the 42ms encompassing the interspike interval for the untreated and ranolazine traces shown in B. Each point represents the value for a single action potential during the 2s of 20 Hz stimulation. action potential MP = the mean membrane potential during the 8 ms encompassing the action potential, Inter-AP MP = the mean membrane potential during the 42-ms interspike interval. D) Plot of the mean membrane potential during the last 500 ms of stimulation. Each point represents an animal average derived from at least 4 fibers without (n = 5) and with treatment with ranolazine (n = 5). The horizontal line represents the mean. E and F) Plots of the mean membrane potential during the 30^th^ action potential and the 30^th^ interspike interval. * indicates p = 0.005 (t-test), ** p= 0.01 (t-test).

However, as myotonia was greatly reduced by ranolazine (Novak et al., 2015; Hawash et al., 2017), it was possible that elimination of plateau potentials was secondary to a reduction in the number of myotonic action potentials. We thus changed our stimulation protocol to a 2-s train of 3-ms stimulus pulses delivered at 20 Hz. This firing rate and duration of firing mimics the duration and rate of firing during runs of myotonia (Hawash et al., 2017). For trains of stimuli, the amplitude of 3-ms pulses of current were first adjusted to find the lowest current required to elicit an action potential. The current was then increased by 10 nA prior to delivering a train of stimuli. 2s of 20 Hz stimulation triggered plateau potentials in 46/55 fibers (n = 5 mice, Fig 5B). Out of the 46 fibers with plateau potentials triggered by 20 Hz trains of stimuli, 11 fibers entered plateau potentials with limited (3 or less myotonic APs) or no myotonia. When 40 μM ranolazine was applied, plateau potentials developed in 0/68 fibers (n = 5 mice, p < .01 vs untreated, Fig 5B). These data suggest that ranolazine is not eliminating plateau potentials via a secondary effect of prevention of repetitive firing.

As shown in Fig. 3C and D, the mean membrane potential at the end of a run of myotonia correlated with whether myotonia terminated in a plateau potential or with repolarization. Thus, we tested if ranolazine prevented plateau potentials by decreasing the mean membrane potential prior to development of plateau potentials. Untreated fibers with additional action potentials and plateau potentials occurring during the 2-s stimulation (myotonia) were excluded from analysis to ensure that the presence of myotonia or plateau potentials did not account for the difference in mean membrane potential. As all untreated fibers initially had plateau potentials or myotonia during repetitive stimulation, the analysis was not possible. To address this issue, the warm-up phenomenon was induced using 8 Hz 10s trains, which lessens myotonia due to slow inactivation of Na channels (Novak et al., 2015). It must thus be noted that the untreated muscle was not at baseline but had already undergone some Na channel inactivation. No induction of warm-up was necessary following treatment with ranolazine, as no fibers had myotonia or plateau potentials. In the absence of ranolazine, the mean membrane potential during the 500ms prior to the plateau potential was −49.6 ± 2.5 mV (n = 5 muscles, 28 fibers). In the presence of 40 μM ranolazine, the mean membrane potential was less depolarized (−56.2 ± 2.9 mV, p < .05 vs untreated, n = 5 muscles, Fig 5D). These data suggest that ranolazine prevents plateau potentials by lessening depolarization of the mean membrane potential during the 20 Hz stimulation.

There are two contributors to the mean membrane potential during repetitive stimulation: 1) the membrane potential during action potentials, and 2) the membrane potential during the interspike interval. We analyzed the contribution of each of these to the hyperpolarization caused by ranolazine. By the 30^th^ action potential of the 20 Hz stimulation, action potential duration had increased to close to 8ms. This widening of the spike-form was likely due to failure of the membrane potential to fully repolarize between action potentials, given that depolarization has previously been found to cause widening of action potentials (Renaud and Light, 1992; Yensen et al., 2002; Miranda et al., 2017). We examined the mean membrane potential during the 8ms encompassing the 30^th^ action potential and found a trend (not statistically significant) toward lessening of depolarization following treatment with ranolazine (Fig 5C, E: −28.2 ± 3.4 vs −32.9 ± 3.8 mV, p = 0.07). With 20 Hz stimulation, there is an action potential every 50ms. After taking the mean membrane potential for the 8ms encompassing the 30th action potential, there remained 42ms in which there was no action potential. The membrane potential for this 42ms interspike interval before the 30^th^ action potential was less depolarized following treatment with ranolazine (Fig 5C, F: −56.0 ± 2.3 vs −61.7 ± 3.0 mV, p < .05). As the interspike interval accounts for 84% (42/50ms) of the time between spikes during 20 Hz stimulation, hyperpolarization of this interval is largely responsible for hyperpolarization of the mean membrane potential.

### Ranolazine prevents transient weakness *in vivo*

The finding that ranolazine eliminated plateau potentials allowed us to explore whether plateau potentials are the mechanism underlying transient weakness *in vivo*. We recorded triceps surae force in 5 ClC^adr^ myotonic mice before and 45 minutes after intraperitoneal (i.p.) injection of 50 mg/kg of ranolazine. As shown previously(Novak et al., 2015), treatment with ranolazine decreased myotonia such that muscle was able to more rapidly relax following termination of stimulation (Fig 6A). In addition to lessening myotonia, treatment with ranolazine eliminated transient weakness in all 5 mice (Fig 6A and B, p < .01 vs untreated at 16s). The elimination of both plateau potentials and transient weakness by ranolazine supports the hypothesis that plateau potentials are the mechanism underlying transient weakness *in vivo*.

**Fig 6:**
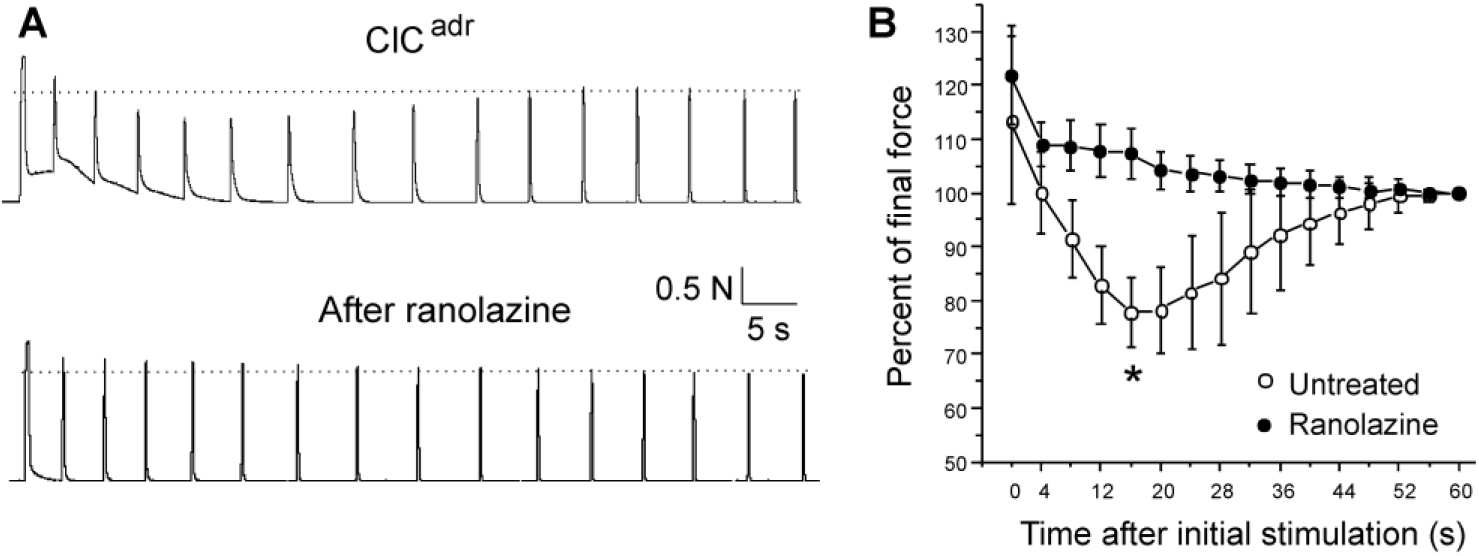
Ranolazine eliminates transient weakness *in vivo.* A) Shown are force traces from a triceps surae muscle group before and 45 minutes after i.p. injection of 50 mg/kg ranolazine. B) Shown is the mean normalized force in 5 ClC^adr^ mice before and after injection of ranolazine. p = .001 for the difference in force 16s (*) after the initial stimulation (paired t-test, 95% confidence interval 70-86 vs 102-113).

## Discussion

Motor dysfunction in recessive myotonia (Becker disease) involves both muscle stiffness and transient weakness. Using intracellular recordings from a mouse model of myotonia congenita (ClC^adr^), we discovered that, while some runs of myotonia resolved with repolarization, others terminated with a plateau potential; that is, depolarization to a membrane potential between −30 and −45 mV, lasting up to 100s. There was gradual repolarization during plateau potentials until the membrane potential reached −45 mV, at which point a sudden repolarization to the resting membrane potential occurred. During plateau potentials, muscle fibers were inexcitable. Studies of genetic and pharmacologic mouse models suggest both current through voltage-activated Ca_v_1.1 Ca^2+^ channels and sodium persistent inward current (NaPIC) may contribute to plateau potentials. Na^+^ persistent inward current (NaPIC) through Na_v_1.4 channels is the key trigger of plateau potentials and current through Ca_v_1.1 Ca^2+^ channels contributes to sustaining plateau potentials. Blocking NaPIC with ranolazine eliminated both plateau potentials *in vitro* and transient weakness *in vivo*. Our results suggest plateau potentials are the mechanism underlying transient weakness in Becker disease.

### Rapid transition between hyperexcitability and inexcitability in Becker Disease

Our data suggest that muscle from a mouse model of Becker disease undergoes a rapid transition between the states of hyperexcitability (myotonia) and inexcitability (due to plateau potentials), shown in Fig 7. Repeated firing of action potentials during voluntary contraction triggers myotonia, which often transitions to a depolarization that forms a plateau potential. During plateau potentials, fibers cannot generate action potentials in response to stimulation, providing an explanation for the drop in CMAP amplitude reported in patients (Ricker and Meinck, 1972; Brown, 1974; Aminoff et al., 1977; Deymeer et al., 1998; Drost et al., 2001; Modoni et al., 2011). The reason for the rapid transition between myotonia (hyperexcitability) and plateau potentials (inexcitability) is that both states are caused by depolarization secondary to loss of muscle Cl^-^ current. The difference is one of degree: when depolarization is mild, Na^+^ channels are not inactivated, such that repetitive firing of action potentials is triggered. When depolarization worsens, Na^+^ channels inactivate, such that inexcitability and paralysis ensue. This proposal is similar to the current understanding of hyperkalemic periodic paralysis, in which there is often myotonia at the beginning of attacks (during the initial, mild depolarization) and weakness at the height of an attack (when depolarization is maximal) (Cannon, 2015; Statland et al., 2018).

**Fig 7:**
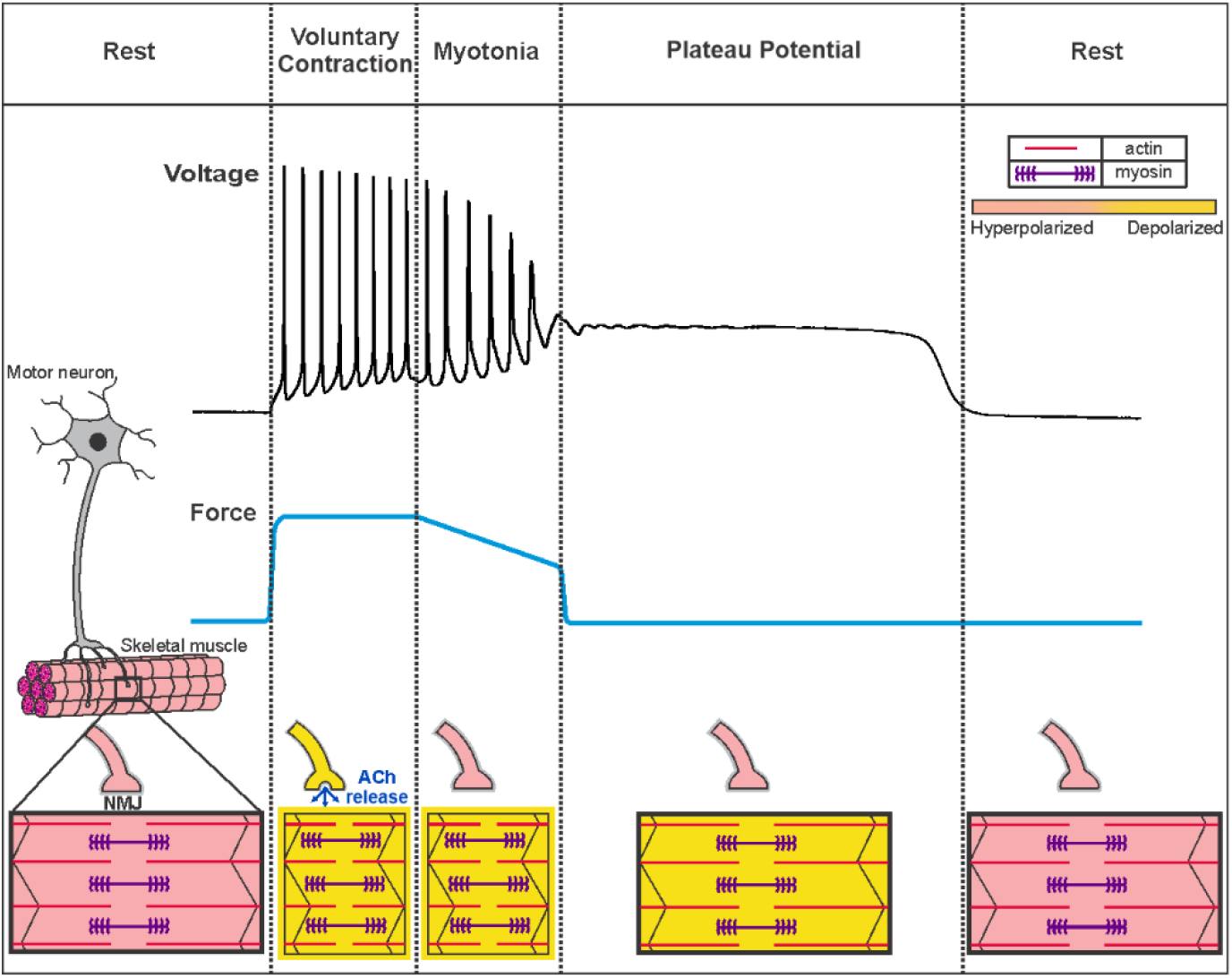
Voluntary contraction in myotonia congenita triggers a sequential progression through states of hyperexcitability and inexcitability. Shown on the left is a motor unit consisting of a motor neuron and the muscle fibers it innervates. At rest, both the motor neuron and muscle fibers are hyperpolarized and muscle is relaxed. To initiate voluntary contraction, the motor neuron fires repeated action potentials, which activate the neuromuscular junction to trigger repeated firing of action potentials in muscle (indicated by a yellow outline around the fiber). With the rapid firing of muscle action potentials, there is sustained contraction and force production (blue line). At the end of voluntary contraction, the motor neuron stops firing and the neuromuscular junction (NMJ) repolarizes, but in myotonic muscle there is continued involuntary firing of action potentials, which slows relaxation of muscle. When myotonia is terminated by transition into a plateau potential, muscle remains depolarized but cannot fire action potentials (black outline of the fiber), such that muscle is paralyzed. Finally, there is sudden repolarization and return of muscle to the resting state. ACh = acetylcholine.

### Ion channels contributing to generation and maintenance of plateau potentials

Involvement of voltage-gated channels in the generation of plateau potentials is suggested by the strong correlation between membrane potential and the initiation and termination of plateau potentials. In runs of myotonia that terminated in a plateau potential, the mean membrane potentials averaged to include both action potentials and interspike intervals prior to development of a plateau potential was −40 mV; whereas, within the same fibers, runs of myotonia terminating with repolarization had a mean membrane potential prior to repolarization of −49 mV. The midpoint of the difference between these values is close to −44 mV. In ClC^adr^ muscle, the mean membrane potential prior to termination of plateau potentials was −43 mV, and in 9-AC treated muscle it was −45 mV. Thus, a mean membrane potential of −44 mV appears to predict both entry into, as well as termination of, plateau potentials.

Both Ca_v_1.1 and Na_v_1.4 are voltage-gated channels that could depolarize muscle to potentials achieved during plateau potentials. In spinal motor neurons, CaV1.3 channels play a central role in generation of plateau potentials that last many seconds (Alaburda et al., 2002; Heckman and Enoka, 2012; Hounsgaard, 2017). Here, we show that the development of plateau potentials was not affected in *nc*DHPR muscle that lack Ca^2+^ current through Ca_v_1.1. However, there was a reduction in duration of plateau potentials in *nc*DHPR muscle. These data suggest that Ca^2+^ flux through Ca_v_1.1 channels contributes to sustaining plateau potentials. This was a surprise, as the membrane potential during plateau potentials is more negative than voltages at which there is significant current through Ca_v_1.1 channels (Garcia and Beam, 1994; Bannister and Beam, 2013). One explanation is that in intact, mature fibers, prolonged depolarization during plateau potentials allows for activation of Ca_v_1.1 at more negative potentials than during the shorter step depolarizations typically used for voltage clamp studies. Our findings suggest that despite Ca^2+^ current through Ca_v_1.1 having no essential role in healthy muscle (Dayal et al., 2017), it may contribute to pathologic depolarization in muscle channelopathies.

Partial block of Na_v_1.4 channels with ranolazine eliminated plateau potentials, suggesting involvement of Na_v_1.4. However, the majority of Na_v_1.4 channels inactivate within ms (fast-inactivating Na_v_1.4 channels), such that they cannot contribute to prolonged depolarization during plateau potentials. There is a Na^+^ persistent inward current (NaPIC) in skeletal muscle that appears to arise from a subset of Na_v_1.4 channels that lack fast inactivation (Patlak and Ortiz, 1986; Gage et al., 1989; Hawash et al., 2017). This subset of Na_v_1.4 channels could play a role in the development of plateau potentials.

Ranolazine has been found to preferentially block NaPIC (El-Bizri et al., 2011; Kahlig et al., 2014). When myotonic muscle was treated with ranolazine to block NaPIC, depolarization of the mean membrane potential during repetitive firing was decreased and plateau potentials were prevented. An important contributor was lessening of depolarization of the membrane potential during the interspike interval. Previously, the only identified contributor to depolarization of the interspike membrane potential was K^+^ build-up in the transverse (t)-tubules (invaginations of the sarcolemma), which depolarizes the K^+^ equilibrium potential (Adrian and Bryant, 1974; Adrian and Marshall, 1976; Wallinga et al., 1999; Fraser et al., 2011). The finding that block of NaPIC lessens depolarization of the interspike membrane potential suggests that NaPIC is activated during this interval.

We hypothesize that activation of NaPIC combines with K^+^ build-up in t-tubules to depolarize muscle to a mean membrane potential of −44 mV during myotonia, such that a plateau potential is triggered. An additional contributor to this depolarization may be the lessening of inward rectifier potassium channel (Kir) conductance with depolarization (Standen and Stanfield, 1980; Struyk and Cannon, 2008). It is unclear whether NaPIC, K^+^ build-up, and decreased Kir conductance can fully account for depolarization during plateau potentials. Based on studies of sustained depolarizations (sometimes termed plateau potentials) in neurons, a family of ion channels that might contribute is the transient receptor potential (TRP) ion channel family (Yan et al., 2009; Phelan et al., 2012). Members of the TRP ion channel family are expressed in skeletal muscle (Brinkmeier, 2011; Gailly, 2012), and we recently found that activation of TRPV4 plays a role in triggering percussion myotonia (Dupont et al., 2020).

It is likely that voltage-gated ion channels promoting repolarization are involved in the sudden termination of plateau potentials. Oscillations in membrane potential often occurred at the beginning or end of plateau potentials. In motor neurons, oscillations in membrane potential are caused by a balance between 2 voltage-gated ion channels: one promoting depolarization and one promoting hyperpolarization (Iglesias et al., 2011; Sciamanna and Wilson, 2011; Nardelli et al., 2017). Kv channels could participate in oscillations occurring during plateau potentials given that Kv conductance in muscle is large and the channels begin to activate at voltages reached during plateau potentials (Beam and Donaldson, 1983; DiFranco et al., 2012). Thus, it may be possible to prevent plateau potentials by either blocking channels promoting depolarization such as NaPIC, or by opening channels promoting repolarization such as Kv channels.

### Blocking NaPIC as therapy for transient weakness

In a mouse model of myotonia congenita, ranolazine prevented both development of plateau potentials *in vitro* and transient weakness *in vivo.* In open label trials of ranolazine in both myotonia congenita and paramyotonia congenita, there were statistically significant reductions in the degree of self-reported weakness (Arnold et al., 2017; Lorusso et al., 2019). Taken together, these data suggest that the clinical benefit of blocking Na^+^ channels in some diseases with myotonia may result, in part, from prevention of transient weakness secondary to development of plateau potentials.

Mutations of Na_v_1.4 responsible for hyperkalemic periodic paralysis increase NaPIC (Cannon et al., 1991; Cannon and Strittmatter, 1993), which appears to play a central role in triggering the depolarization that underlies attacks of transient weakness (Lehmann-Horn et al., 1987; Jurkat-Rott et al., 2010; Cannon, 2015). This suggests that blocking NaPIC should be effective in treating hyperkalemic periodic paralysis. However, treating patients with Na channel blockers that are effective in treating myotonia, such as mexiletine, has previously been found to be ineffective (Ricker et al., 1983; Ricker et al., 1986). The finding that blocking NaPIC with ranolazine lessens depolarization of the interspike membrane potential suggests it might be worth considering a trial of this FDA-approved drug’s efficacy in hyperkalemic periodic paralysis.

### Conclusion

We identified currents contributing to plateau potentials that are responsible for transient weakness in recessive myotonia congenita. We also determined that blocking a current contributing to plateau potentials provided effective amelioration of transient weakness. Currents contributing to plateau potentials are present in wild-type muscle; thus, they might contribute to depolarization in other muscle channelopathies with transient weakness, such as hyper- and hypokalemic periodic paralysis. Identification of channels involved in generation of plateau potentials in skeletal muscle may thus advance understanding of regulation of excitability in both healthy and diseased muscle.

## Materials and Methods

### Mice

All animal procedures were performed in accordance with the policies of the Animal Care and Use Committee of Wright State University and were conducted in accordance with the United States Public Health Service’s Policy on Humane Care and Use of Laboratory Animals.

The genetic mouse model of myotonia congenita used was *Clcn1^adr-mto^/J* (ClC^adr^) mice, which have a homozygous null mutation in the *Clcn1* gene (Jackson Laboratory Stock #000939). The pharmacologic model of Becker disease involved treatment of muscle with 100 μM 9-anthracenecarboxylic acid (9-AC). The mouse model of non-Ca^2+^-conducting Ca_v_1.1 used was *nc*DHPR, carrying point mutation N617D in pore loop II in the *Cacna1S* gene (Dayal et al., 2017).

Genotyping of ClC^adr^ mice was performed as previously described to select heterozygous mice for breeding (Dupont et al., 2019). Otherwise, homozygous myotonic mice were identified by appearance and behavior as previously described (Novak et al., 2015). Unaffected littermates were used as controls. Genotyping for selection of homozygous *nc*DHPR was performed as previously described (Dayal et al., 2017). Both male and female mice were used from 2 months to 6 months of age. As mice with myotonia have difficulty climbing to reach food, symptomatic mice were supplied with moistened chow paste (Irradiated Rodent Diet; Harlan Teklad 2918) on the floor of the cage.

### Electrophysiology

Current and voltage clamp recordings were performed at 20–22°C.

### Current clamp

Mice were sacrificed using CO_2_ inhalation followed by cervical dislocation, and both extensor digitorum longus (EDL) muscles were dissected out tendon-to-tendon. Muscles were maintained and recorded at 22°C within 6 hours of sacrifice. The recording chamber was continuously perfused with Ringer solution containing (in mM) NaCl, 118; KCl, 3.5; CaCl_2_, 1.5; MgSO_4_, 0.7; NaHCO_3_, 26.2; NaH_2_PO_4_, 1.7; glucose, 5.5 (pH 7.3-7.4 at 20-22°C), and equilibrated with 95% O_2_ and 5% CO_2_.

Intracellular recordings were performed as previously described (Novak et al., 2015; Hawash et al., 2017; Dupont et al., 2019). Briefly, muscles were loaded with 50 μM N-benzyl-p-toluenesulfonamide (BTS, Tokyo Chemical Industry) for at least 30 minutes prior to recording to prevent contraction. BTS was dissolved in DMSO and added to the perfusate prior to bubbling with 95% O_2_ and 5% CO_2_, as we have found that BTS is more soluble at a basic pH. Prior to recording, muscles were stained for 3 minutes with 10μM 4-(4-diethylaminostyrl)-N-methylpyridinium iodide (4-Di-2ASP, Molecular Probes) to allow imaging muscle with an upright epifluorescence microscope (Leica DMR, Bannockburn, IL).

Micro-electrodes were filled with 3M KCl solution containing 1 mM sulforhodamine to visualize the electrodes with epifluorescence. Resistances were between 15 and 30 MΩ, and capacitance compensation was optimized prior to recording. Fibers with resting potentials more depolarized than –74 mV were excluded from analysis.

In cases where there was failure of action potentials during trains of stimulation, we did not determine whether the failure was due to the presence of an absolute or relative refractory period by altering current injection during the train of stimuli.

### Voltage clamp

*Flexor digitorum brevis* (FDB) and *interosseous* (IO) muscle fibers were isolated as previously described (Waters et al., 2013; Hawash et al., 2017). Briefly, muscles were surgically removed and enzymatically dissociated at 37 °C under mild agitation for ~1 h using 1,000 U/mL of collagenase type IV (Worthington Biochemical). Mechanical dissociation was completed using mild trituration in buffer with no collagenase. The fibers were allowed to recover at 20-22 °C for 1 hour before being used for electrical measurements.

Both the current-passing and voltage-sensing electrodes were filled with internal solution (see below) and had resistances of ~15 MΩ. After impalement, 10 minutes of hyperpolarizing current injection was allowed for equilibration of the electrode solution. Data were acquired at 20 kHz and low-pass filtered with the internal Axoclamp 900A filters at 1 kHz. The voltage clamp command signal was low-pass filtered with an external Warner LFP-8 at 1 kHz.

Internal solution (in mM) was as follows: 75 aspartate, 30 EGTA, 15 Ca(OH)_2_, 5 MgCl_2_, 5 ATP di-Na, 5 phosphocreatine di-Na, 5 glutathione, 20 MOPS, and pH 7.2 with CsOH. Extracellular solution (in mM) was as follows: 144 NaCl, 4 CsCl, 1.2 CaCl_2_, 0.6 MgCl_2_, 5 glucose, 1 NaH_2_PO_4_, 10 MOPS, 0.05 BaCl_2_, 0.1 9-AC, 0.001 TTX, 0.01 Ouabain, 0.1 3,4-diaminopyridine (3,4-DAP), and pH 7.4 with NaOH.

### Drugs and doses (when used, in mM)

0.05 BTS/N-benzyl-*p*-toluenesulfonamide (TCI America, Portand OR)
0.1 9-AC/9-anthracenecarboxylic acid (Tocris Bioscience, Minneapolis, MN)
0.05 Ranolazine (Sigma-Aldrich, St. Louis, MO)
0.001 TTX/Tetrodotoxin (Alomone Labs, Jerusalem, Israel)
0.1 3,4-DAP/3,4-Diaminopyridine (Sigma-Aldrich, St. Louis, MO)
0.01 Ouabain (Sigma-Aldrich, St. Louis, MO)

### In situ muscle force recording

*In situ* muscle force recordings were performed as previously described (Dupont et al., 2019; Wang et al., 2020). Briefly, mice were anesthetized via isoflurane inhalation; then the distal tendon of the triceps surae muscles was attached to a force transduction motor and the sciatic nerve was stimulated while isometric muscle force generation was measured. The sciatic nerve was stimulated with constant current injection. The amplitude of the current pulse was adjusted to 150% of the current required to trigger a single action potential. To induce myotonia, 45 pulses of 1ms duration were delivered at 100 Hz. To follow development of transient weakness, 15 pulses delivered at 100 Hz were delivered every 4s. Muscle temperature was monitored with a laser probe and maintained between 29°C and 31°C with a heat lamp. The muscle was kept moist by applying mineral oil. Ranolazine was administered via intraperitoneal (i.p.) injection at a dose of 50 mg/kg dissolved in water. The typical volume of water injected was 100 μl.

### Statistics

Sample size was determined by past practice, where we have found an n of 5 muscles from 5 different mice, studied on different days, yields statistically significant differences in muscle action potential properties. At least 5 muscle fibers were recorded from in each muscle. However, for studies of plateau potentials in ClC^adr^ mice we obtained recordings with adequate preservation of resting membrane potential in only a subset of fibers so fewer fibers per muscle were included in the final analysis. This was due to the prolonged duration of plateau potentials, which required maintaining prolonged impalement of individual fibers with two electrodes. No outlier data points were excluded. Intracellular recording data from different mice were analyzed using nested analysis of variance with n as the number of mice, with data presented as mean ± SD. *p* < 0.05 was considered to be significant. The numbers of animals and fibers used are described in the corresponding figure legends and text. For parameters that were not normally distributed, such as duration of the plateau potential and slope of the repolarization, differences between two data sets were analyzed after applying a natural log transformation, which yielded normally distributed data.

For comparisons of data within individual muscle fibers (mean membrane potential and mean firing rate for runs of myotonia, without and with plateau potentials), the paired student’s t-test was used with n as the number of fibers. For force recording comparisons before and after ranolazine treatment, the paired student’s t-test was used with n as the number of mice. For comparisons of force recordings between myotonic mice and unaffected littermates, a 2-sample student’s t-test was used.

## Acknowledgements

This work was supported by NIH grants AR074985 (M.M.R.), and MDA grant 602459 (M.M.R.).

## Author contributions

JM, KD, CD, AH, KN and AK all acquired, analyzed data, and made figures. MG, AD, AV and MR drafted and revised the manuscript.

